# The methodological foundations of lesion network mapping remain sound

**DOI:** 10.64898/2026.02.24.707529

**Authors:** Shan Siddiqi, Andreas Horn, Frederic L.W.V.J. Schaper, Sanaz Khosravani, Alexander Li Cohen, Juho Joutsa, John D. Rolston, Michael A. Ferguson, Samuel B. Snider, Anderson M. Winkler, Harith Akram, Stephen M. Smith, Thomas E. Nichols, Karl Friston, Aaron D. Boes, Michael D. Fox

**Author notes:** Corresponding authors: Michael D. Fox, MD, PhD, Center for Brain Circuit Therapeutics, 60 Fenwood Rd, Boston MA 02115, Shan H. Siddiqi, MD, Dauten Behavioral Health Institute, 710 N Lake Shore Dr, Chicago, IL 60611.

## Abstract

Lesion network mapping (LNM) and related techniques have been used in over 200 studies, primarily to test whether anatomically distributed lesions that cause the same symptom fall within a common brain network. A recent article^1^ challenges the specificity and validity of this technique, suggesting that lesion network maps primarily reflect intrinsic properties of the normative connectome rather than lesion–symptom relationships. However, the data and procedures in van den Heuvel et al. do not reflect those used in most LNM studies. Further, the main conclusions were based on similarity between maps, but similarity does not imply the absence of meaningful differences. In contrast, LNM provides evidence for meaningful differences using specificity testing. Exemplary analyses of 1090 lesion locations from 34 prior LNM studies do not support van den Heuvel’s concerns and confirm the lesion-deficit specificity of LNM. While we encourage further methodological investigation, the analyses of van den Heuvel et al. do not invalidate prior LNM findings or future applications.

---

We thank van den Heuvel et al. for their methodological investigation into lesion network mapping and related techniques. Given the increasing uptake of this technique and its potential for clinical translation, efforts to continuously evaluate it are welcome. Prior LNM studies tend to follow a similar methodological approach (Figure 1A). Step 1 includes sensitivity testing for common network connections across distributed focal lesions causing the same symptom, sign, syndrome, or condition. Step 2 includes specificity testing, where the set of lesions causing a particular symptom is compared to a set of control lesions causing different symptoms. This step ensures the lesion-deficit mapping has face validity and is specific to the symptom in question.

**Figure 1:**
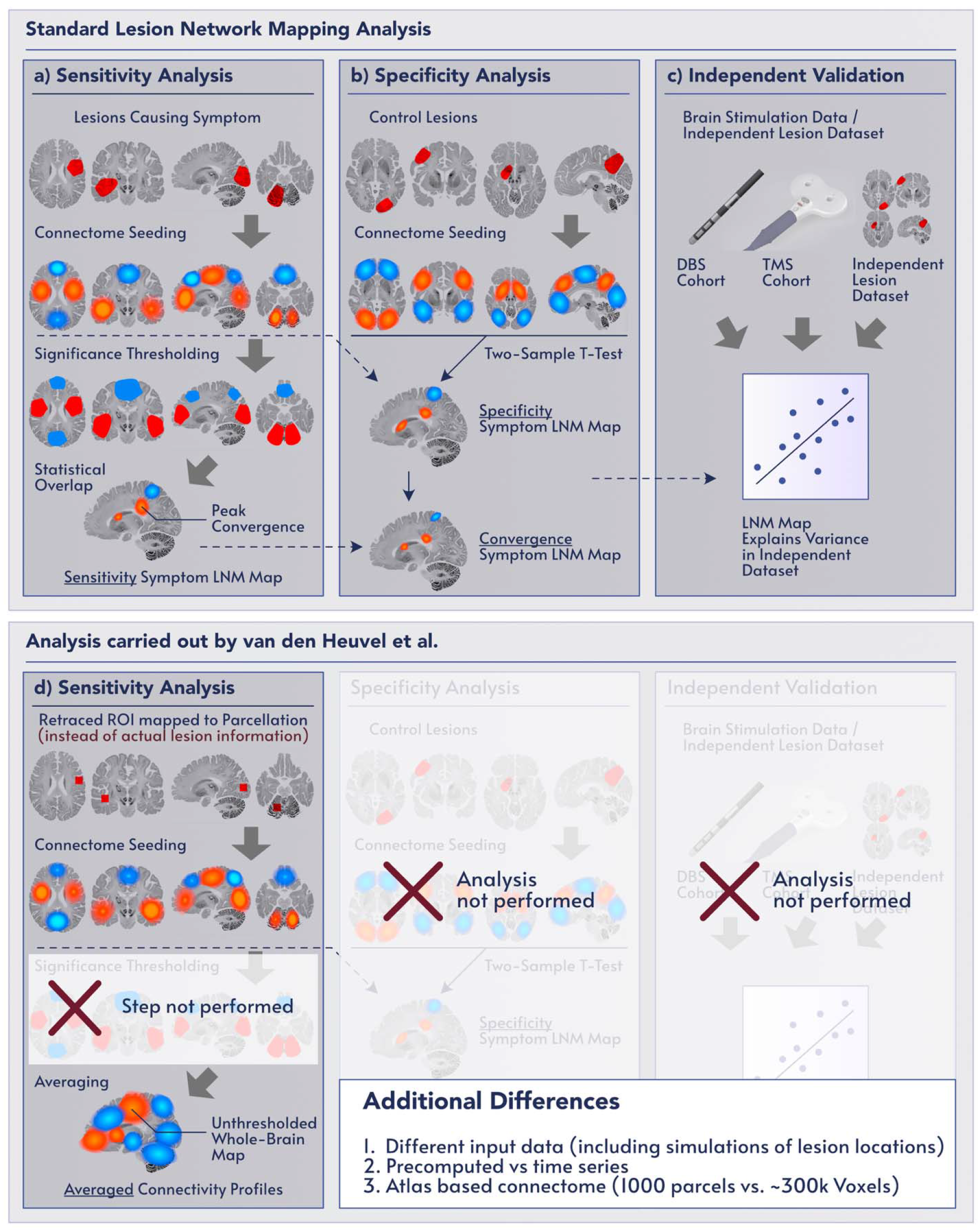
Standard lesion network mapping methods vs. the modified method used by van den Heuvel et al. The top panel outlines typical methodology used by most published lesion network mapping publications. The bottom panel summarizes the modified method used by Van den Heuvel et al. for most analyses. Key differences between the two approaches include the use of alternate input data, map averaging rather than statistical sensitivity testing, omission of specificity testing, omission of conjunction testing, and omission of validation against independent data modalities.

In step 3, LNM studies often establish construct validity using independent lesion datasets, independent modalities such as brain stimulation, or different connectomes that do not rely on functional connectivity.^2–6^ For example, lesions relieving tremor show peak connectivity to the same thalamic voxels in which implanted deep brain stimulation (DBS) electrodes are proven to relieve tremor^2^ while lesions causing depression are connected to the same brain network as transcranial magnetic stimulation (TMS) sites that relieve depression^4^. Furthermore, a recent head-to-head randomized trial found that targeting different LNM-derived networks with TMS specifically improved the corresponding psychiatric symptoms^7^.

Van den Heuvel et al. challenge this body of work by noting that lesion network maps 1) can appear similar across different symptoms and 2) reflect properties of the connectome, namely, its degree. These observations are perfectly sensible. Indeed, the rationale for LNM is to characterize lesion-deficit mappings under the constraints offered by the connectome. The resulting lesion network maps may appear similar. The relevant question is whether any component of these maps is specific to the symptom caused by the lesion. This is the reason specificity testing is routinely included in LNM studies (Figure 1). We also agree with some other points raised by van den Heuvel et al. including the value of quantitatively comparing LNM results across published studies and calculating the false positive rate of the LNM technique to derive null models.

However, we disagree with the assertion that lesion network maps contain “no substantive information other than unspecific signal”, as this claim contradicts the results of multiple prior LNM papers. To investigate this discrepancy, we re-assessed our database from published LNM studies including 1090 lesion locations causing 34 different symptoms (Table S1). Using van den Heuvel et al.’s preferred metric of spatial correlation between unthresholded lesion network maps, we tested three assertions:

## Assertion 1: LNM results are not symptom-specific

Our data do not support this assertion, as the spatial correlation between lesion networks from patients with the same symptom was significantly higher than between those with different symptoms (median spatial r= 0.44 vs 0.09; p < 0.0001).

## Assertion 2: LNM results all converge to degree maps

Our data do not support this assertion, as the spatial correlation between lesion networks from patients with the same symptom was significantly higher than the spatial correlation between these networks and the degree map (median spatial r = 0.44 vs 0.16, p< 0.0001).

## Assertion 3: LNM is prone to false positive results because of flawed specificity testing

Our data do not support this assertion, as we did not identify a single false positive in 1000 iterations when we compared 50 lesions randomly extracted from our database to the remaining 1040 lesions, using the same thresholds applied by van den Heuvel et al. (sensitivity 75%, specificity t >10). False positives (4.6% of permutations) only began to emerge when thresholds were dropped below the limits used in most prior LNM studies (e.g. sensitivity 75%, specificity t = 3.0). We also tested whether the specificity maps generated from these random comparisons would match each other and the topography of the degree map. We found that specificity maps were different from each other (average spatial r [SD] = -0.0003 [0.3490]) and from the degree map (average spatial r [SD] = -0.0026 [0.3493]).

Taken together, these results suggest that LNM 1) shows symptom specificity across studies, 2) produces results that do not converge on a single map, and 3) controls for false positives with specificity testing. We next sought to understand why the findings of van den Heuvel et al. differed from prior LNM studies and the above analyses. We highlight four main factors that may have contributed to this discrepancy:

First, the main conclusions of van den Heuvel et al. are based on a simplified version of LNM that differs from the method used in prior papers (Figure 1). While there are many differences, the most important may be the omission of specificity testing designed to preclude non-specific patterns of connectivity, including the degree of the connectome. Similar to how only brain activations associated with a specific task (versus a control task) are considered part of an activation network in task-based fMRI^8^, only connections that survive tests for specificity tests are usually considered part of a lesion network. LNM findings that survive this statistical testing, especially the peak findings, can differ meaningfully across symptoms and studies, even when the unthresholded whole-brain maps appear similar (Figure 2). Van den Heuvel et al. consider these issues and justify their deviations from standard LNM methods based on a simulation reported in the supplement of their paper (S15). They report a high false positive rate despite applying statistical sensitivity and specificity testing, leading the authors to conclude that these steps do not add value and can be omitted. Using real rather than simulated lesion data, we obtained a very low false positive rate (see above). These seemingly contradictory results stem from assumptions in van den Heuvel’s simulations. When they assumed low overlap in the lesion locations, they obtained a low false positive rate, consistent with our randomly selected real lesion data. Only when they assumed high overlap in lesion locations did they obtain high false positive rate. However, high overlap in lesion locations is not a property of randomly selected lesions, but it is a property of lesions selected based on causing a common symptom. As such, these simulated “false positives” could be interpreted as “true positives” that accurately reflect the properties of the input data. In short, van den Heuvel’s simulation did not produce significant findings with random lesion locations, but did produce significant findings with non-random lesion locations, which could be interpreted as supporting the importance of specificity testing in LNM rather than justifying its omission.

**Figure 2:**
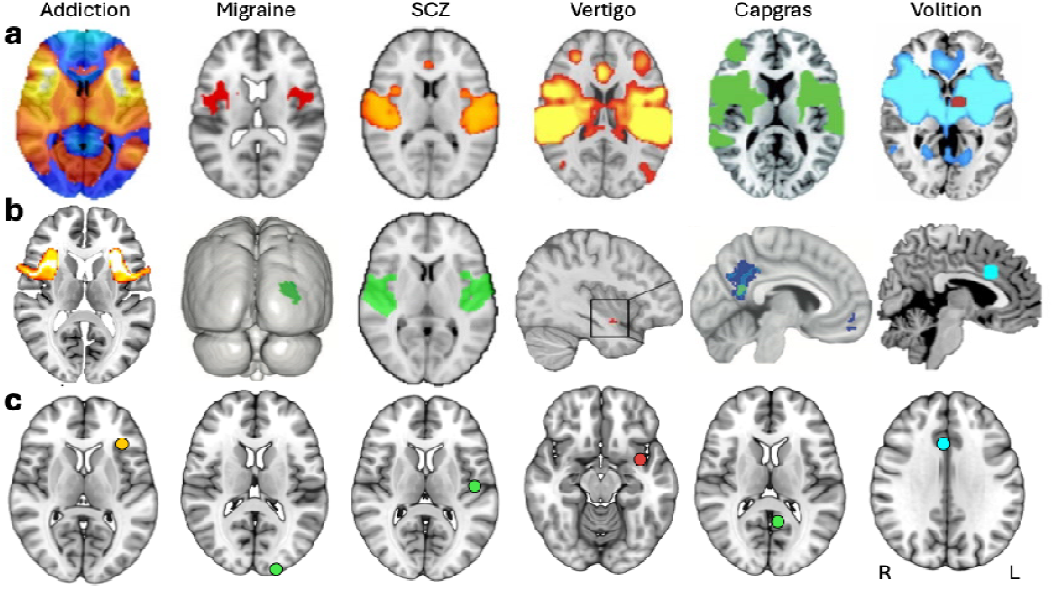
Network mapping results from six published studies selected by van den Heuvel et al. to illustrate a lack of specificity. A) Figure 1a from van den Heuvel et al., with images presumably selected to illustrate the similarity of network mapping results across studies. B) Images extracted from the same articles, showing the peak finding from each study that was robust to statistical sensitivity and specificity testing. C) Spheres at the peak coordinates reported in each study, displayed on axial slices to better facilitate comparison between studies. Abbreviations: SCZ, schizophrenia; R, right; L, left.

Second, van den Heuvel et al. only tested for similarity between maps, not for equivalence or lack of differences. They report that almost all maps examined were more similar than expected by chance based on spin-test^9^ and BrainSMASH^10^, two methods that attempt to account for spatial autocorrelation by modelling a null distribution. However, these methods have been criticized since they make assumptions about the distribution of the input data, do not account for geometric distortions^11^, and can inflate the statistical significance of spatial similarity metrics^12,13^. An alternative method used in many prior LNM studies assesses similarity using permutation testing of the raw input data, which creates a null distribution of spatial correlations by randomly shuffling each patient’s clinical symptom with a different patient’s network map^4,14^. This method makes fewer assumptions, avoids significance inflation when testing for similarity between maps associated with different symptoms, and can also test for differences. To provide just one example, we compared two lesion network maps (for addiction remission and MS-related depression), which were reported to be significantly similar to one another by van den Heuvel et al. (r=0.56, p<0.001), but were not more similar than chance when permuting the raw input data (r=0.56, p=0.29). Moreover, comparing the connectivity of lesions associated with addiction remission lesions versus those associated with MS-related depression using permutation-based specificity testing identified significant differences, despite the above similarity (p_FWE_<0.05).

Third, the authors interpret spatial similarity across LNM results as problematic. Our interpretation of this finding differs. The authors report an average absolute spatial correlation of |r|=0.4 between maps, resulting in r^2^=16% shared variance. Based our analyses and those of independent groups^13^ this may be an over-estimate, but even 16% shared variance does not preclude the existence of biologically meaningful or statistically significant differences.. When one tests for differences—as in numerous prior LNM papers and the above analysis of 1,090 lesion locations —one finds significant differences in associated networks among different symptoms. Spatial similarity between maps also does not invalidate the LNM method. By analogy, human genomes contain >99.9% shared variance^15^, but this does not call into question the methodological foundations of DNA sequencing or the ability of DNA analyses to detect differences between humans. Finally, spatial similarity across findings is not unique to LNM but is present in studies of functional brain activation^8,16,17^, brain metabolism^18^, and regional brain volume^19^. Anatomical similarity may also be interpreted as evidence of shared anatomical connectivity^20^, shared genetics^21^, symptom comorbidity^22,23^, or domain-general neuroanatomy^24^, rather than a statistical inconvenience or concern.

Fourth, some conclusions in van den Heuvel et al. may stem from their “compressed” mathematical formulation of LNM and assumptions regarding the constituent lesion data. When analyzing their model, they make a key assumption that there is “minimal spatial overlap between lesions.” Uniform random sampling of non-overlapping lesions will by necessity sample the entire normative connectome *C*, with averaged results converging on the row-summation vector of *C* (the degree map). We fully agree with this straightforward mathematical demonstration, assuming the connectome is sampled randomly. However, actual lesions causing specific symptoms violate these assumptions, as they do overlap and their spatial distributions are not random. For example, lesions causing amnesia will repeatedly ‘sample’ the hippocampus and connected brain regions but are less likely to sample the motor cortex^25^.The authors acknowledge that non-randomly sampled lesions will not converge to the degree map, but instead will sample a subset of rows of *C*, “primarily reflecting the inherent FC pattern of the underlying seed region(s).” This is indeed the goal of LNM, to identify the subset of *C* that reflects the inherent connectivity pattern of different lesion locations causing a specific symptom and test whether any connections within these patterns are sensitive *and* specific to the symptom being investigated.

In summary, the authors are commended for their interest in LNM and for motivating this discussion. There may be examples in the LNM literature or specific data types where the concerns raised by van den Heuvel et al. apply, and we would like to encourage LNM authors to look for evidence of these concerns in their data. We would also like to encourage further methodological investigation, especially studies that can provide testable hypotheses for improvement and refinement of the method. In pursuit of this goal, a methodological best practices paper for LNM is under preparation by us. In the meantime, we recommend readers to use the data and methods of previously published LNM papers, not the data and methods presented by van den Heuvel et al., as these are not equivalent. Finally, we highlight that the results presented by van den Heuvel et al. do not undermine the methodological foundations of the lesion network mapping method.

## Acknowledgements

S.S.: NIH (K23MH121657, R01MH136248, R01MH140916, R01MH113929), BrainsWay investigator-initiated grant, the Once Upon a Time Foundation, the Milliken Institute/BD^2^, the Sidney R. Baer Foundation, the Dauten Family Foundation, and the Stahl Family

A.H. was supported by the Schilling Foundation, the German Research Foundation (Deutsche Forschungsgemeinschaft, CRC-1451, 431549029 and CRC-1270 ELAINE, 3–299150580).

J.J. was supported by the Research Council of Finland, Finnish Medical Foundation, Sigrid Juselius Foundation, Päivikki and Sakari Sohlberg Foundation, Signe and Ane Gyllenberg Foundation, Finnish Parkinson Foundation, Turku University Hospital (VTR funds), and University of Turku (private donation).

A.L.C. was supported by the Simons Foundation Autism Research Initiative and the TSC Alliance

M.D.F. was supported by grants from the NIH (R01MH113929, R21MH126271, R21NS123813, R01NS127892, R01MH130666, UM1NS132358), the Kaye Family Research Endowment, the Ellison / Baszucki Family Foundation, the Once Upon a Time Foundation, The Milliken Institute / BD^2^, the Manley Family, Donna and Tom May, and Chuck and Kerri Bean.

F.L.W.V.J.S was supported by the National Institutes of Health (R01NS127892) and American Epilepsy Society (846534).

A.D.B was supported from NIH grants R01NS119896, R01NS062820, R01MH139650, R01MH132074

J.D.R. was supported by the National Institutes of Health (R01NS136297), the Once Upon a Time Foundation, and the Milliken Institute / BD^2^.

A.M.W. receives support from the National Institutes of Health (R01MH139547, U54HG013247, and R01MH138425).

## Author contributions

Conceptualization and initial draft: SHS, MDF, AH, and ADB

Data analysis: SHS, FLWVJS, SK, MAF

Independent consultants: KF, TEN, AMW, JDR, SMS

Preparation of figures: AH, FLWVJS, MAF

Writing and revisions: All authors

