## Supplemental methods and table for "The methodological foundations of lesion network mapping remain sound"

To test three key assertions in van den Heuvel’s critique, we re-analyzed data on 1090 lesions across prior studies of lesion network mapping (Supplemental Table 1). As in the original published studies, each lesion was treated as a seed to estimate whole-brain resting-state functional connectivity using a normative connectome database. The methods for these three tests are as follows:

1. Are LNM results symptom-specific?

We classified the 1090 lesions according to the symptom, syndrome, or disorder (henceforth referred to as “syndrome” for simplicity) associated with that lesion. Of note, we identified 47 lesions that were classified in both the “amnesia” category and the “psychosis” category, as they caused both syndromes. These lesions were removed from the psychosis dataset (reducing psychosis sample size from 153 to 106) to avoid overlap.

Next, we computed spatial correlation of every lesion’s connectivity profile to every other lesion’s connectivity profile both within syndrome and between syndromes. This yielded a matrix of spatial correlations for every syndrome compared to itself, as well as another matrix of spatial correlations for every syndrome compared to the other syndromes. To avoid overpowering the analysis by repeatedly sampling the same lesion, we averaged the within-syndrome correlations and the between-syndrome correlations for each lesion, yielding a single mean within-syndrome correlation and a single mean between-syndrome correlation for each lesion. We then compared the within-syndrome value to the between-syndrome value using a paired t-test. We hypothesized that within-syndrome correlations would exceed between-syndrome correlations.

2. Do LNM results all converge to the degree map?

We downloaded the degree sum map of the connectome from the Github repository accompanying van den Heuvel et al.’s critique (https://github.com/dutchconnectomelab/lesionnetworkmapping/). We compared all 1090 lesions to the degree sum map using spatial correlations. These spatial correlations were compared to the above within-syndrome correlations using a paired t-test. We hypothesized that within-syndrome correlations would exceed lesion correlations with the degree sum map.

3. Identifying false positive rate

We randomly extracted 50 lesions from our database and compared them to the remaining 1040 lesions using voxel-wise two-sample t-test. We repeated this analysis with 1000 different iterations of random lesion selection. In each case, we thresholded the lesions using the same thresholds applied by van den Heuvel et al. (specificity t>10, sensitivity 75%). We hypothesized that less than 5% of random iterations would yield false positive results. As a secondary analysis, we also progressively decreased the specificity threshold until achieving a false positive rate near 5%.

We also used spatial correlations to test whether these randomly-sampled maps would resemble each other and the degree sum map.

**N = 1090 non-overlapping lesions**

Table 1. Clinical syndromes and corresponding lesion cohorts included in the analysis.

*From the 153 reported lesions, 106 were selected after excluding those overlapping with the hallucinations category.

| **Clinical Syndromes** | | **Sample Size** | **Reference** |
| --- | --- | --- | --- |
| 1 | Addiction Remission | 34 | Joutsa et al. (2022) |
| 2 | Akinetic Mutism | 28 | Darby et al. (2018) |
| 3 | Alice in Wonderland | 37 | Friedrich et al. (2024) |
| 4 | Alien Limb | 50 | Darby et al. (2018) |
| 5 | Amnesia | 53 | Ferguson et al. (2019) |
| 6 | Anton’s Syndrome | 24 | Kletenik et al. (2023) |
| 7 | Aphantasia | 12 | Kutsche et al. (2025) |
| 8 | Aphasia | 12 | Boes et al. (2015) |
| 9 | Asterixis | 30 | Laganiere et al. (2016) |
| 10 | Blindsight | 34 | Kletenik et al. (2022) |
| 11 | Coma | 12 | Fischer et al. (2016) |
| 12 | Confabulation | 25 | Bateman et al. (2024) |
| 13 | Cortical Blindness | 35 | Kletenik et al. (2022) |
| 14 | Criminality | 17 | Darby et al. (2017) |
| 15 | Delusions (Misidentification Syndromes) | 17 | Darby et al. (2017) |
| 16 | Delusions (Other) | 15 | Darby et al. (2017) |
| 17 | Depression | 58 | Padmanabhan et al. (2019) |
| 18 | Dystonia (Cervical) | 25 | Corp et al. (2019) |
| 19 | Freezing of Gate | 14 | Fasano et al. (2017) |
| 20 | Hallucinations (Auditory) | 21 | Kim et al. (2021) |
| 21 | Hallucinations (Visual) | 49 | Kim et al. (2021) |
| 22 | Hallucinations (Mixed) | 19 | Kim et al. (2021) |
| 23 | Hemichorea | 29 | Laganiere et al. (2016) |
| 24 | Holmes Tremor | 36 | Joutsa et al. (2019) |
| 25 | Infantile Spasm | 74 | Cohen et al. (2021) |
| 26 | Loss of Consciousness (Prolonged) | 16 | Snider et al. (2020) |
| 27 | Mania | 56 | Cotovio et al. (2022) |
| 28 | Pain (Central Post-Stroke) | 23 | Kim et al. (2023) |
| 29 | Parkinsonism | 29 | Joutsa et al. (2018) |
| 30 | Prosopagnosia | 44 | Cohen et al. (2019) |
| 31 | Psychosis | 106* | Pines et al. (2025) |
| 32 | Tics | 22 | Ganos & Horn (2023) |
| 33 | Tremor Relief | 11 | Joutsa et al. (2018) |
| 34 | Vertigo | 23 | Li et al. (2023) |
